# Exploring the dynamic ticks-camel-pathogens interaction

**DOI:** 10.1101/2024.05.15.594365

**Authors:** JohnMark O. Makwatta, Paul N. Ndegwa, Florence A. Oyieke, Peter Ahuya, Daniel K. Masiga, Merid N. Getahun

## Abstract

The ability of ticks to interact and adapt to different ecologies and hosts determines their vectorial competence for various pathogens, however ticks-livestock-pathogens interaction studies are limited. With our ticks-hosts-pathogens interface studies, we found 14 species of ticks feeding on various livestock. Ticks showed a strong preference for one-humped camels (*Camelus dromedarius*). The camel nostril was the most preferred predilection site. The most prevalent tick species on camels was *Hyalomma rufipes*. We found two novel *Amblyomma gemma* variants which are distinct both morphologically and genetically from previously described *Amblyomma gemma*. The signature odors from camel breath and body were attractive to *H. rufipes*; demonstrating ticks utilize camel-derived metabolites to find their host. Our research shows that *H. rufipes* and camel hosts have unique and shared pathogens showing *H. rufipes*’ vector and camel’s reservoir host qualities. Our study unravels the dynamic interactions between ticks, pathogens, and camels that all influence the likelihood of pathogen adaptation and transmission dynamics.

**IMPORTANCE:** Ticks are obligatory hematophagous arachnids, serving as vectors for a wide array of pathogens that can be transmitted to animals and humans. The ability of ticks to acquire and transmit various pathogens depends on its attraction to quality reservoir host and the survival of the pathogens in ticks’ gut and other tissues. However, the complex dynamics of tick-pathogens interaction and host-seeking behavior remains understudied. This investigation revealed notable variation in tick preference for domestic animals, camel being the most preferred host. Moreover, our spatial analysis about tick attachment sites showed nostril is the most preferred sites by various tick species. Our epidemiology data showed variation in the pathogens harbored by camel (host) and vector (*H. rufipes*), demonstrating the camel’s efficiency as reservoir host and ticks’ vector competence for various pathogens. With our behavioral experiment using *H. rufipes* and its preferred host’s (camel) breath and body signature odors, we identified novel attractants for *H. rufipes*, thus offering new avenues for combating TBDs. Overall, our study presents novel insights into how multiple factors shape tick-host-pathogens interaction.

## Introduction

As obligatory ectoparasites, most tick species interact and feed on a wide range of hosts except for some monoxenic species^1,2^; elements which are impetus for pathogen transmission. Ticks transmit diverse pathogenic microbial (viruses, bacteria) and parasitic (protozoans, helminths) agents to their vertebrate hosts including humans ^3–5^. The role ticks play in disease transmission depends on tick species dynamics, vectorial capacity, host choice, and whether they feed on efficient reservoir hosts or not. For instance, Ginsberg et al. ^6^, clearly demonstrated how tick host choice between efficient and poor reservoir host determine the distribution of Lyme disease^6^. Unlike most blood-sucking vectors such as mosquitoes and other biting flies, fleas, lice and bugs, most ticks have a rather unique life cycle involving multiple stages and hosts ultimately resulting in complex tick-hosts-pathogens interaction, adaptation, and epidemiological patterns of pathogen acquisition and transmission^1,7^. Under natural conditions, an animal may be infested with several ticks of different species^8^. The aggregation of ticks on a given host may be due to various reasons, for instance, host infection with various pathogens increases the attractivity of the host to ticks^9,10^. Furthermore, tick infection alters sites of attachment^2^ that initiate ticks’ movement and exposure to new host-pathogen interaction.

The interactions between the tick and its host-pathogen system and with other transmission cycles, such as abiotic factors, are key features that determine the transmission dynamics and distribution of tick-borne diseases^6^. The evaluation of such interactions is a complex but necessary preliminary step in assessing disease transmission risk. Several studies have addressed the importance of examining the complete community of hosts in a territory to estimate their relative contribution in supporting the various tick species present and the pathogens the ticks transmit ^1,7^’^11–14^. The maintenance of tick-borne pathogens within natural reservoirs is intricately influenced by the attraction of ticks to their host animals ^1,7^. Conversely, studies on the diversity, abundance, ticks’ preferential attraction to various co-herded livestock, tick-host-chemical communication, which are the driving factors for tick-borne pathogen transmission, and tick evolutionary adaptation remain understudied. Furthermore, the elucidation of tick-host-pathogen interactions complementary to understanding tick-borne disease transmission dynamics is an emerging field that holds immense potential for the development of innovative strategies aimed at controlling ticks and curtailing the spread of tick-borne diseases.

The objective of our study was to investigate the complex interactions between ticks, hosts, and pathogens by comparing tick infestations in various domesticated livestock hosts. Additionally, we sought to analyze the network of pathogens and examine the chemical communication between ticks and camels in arid and semi-arid regions. *H. rufipes* exhibited a distinct preference for camels over other livestock when it comes to tick-host interactions. Evidently, scents originating from camels were attractive to *H. rufipes*.

## Results

### Prevalence of ticks on camels and co-herding livestock

The tick prevalence in camels was high as compared to the other co-herding livestock, followed by cattle (*P* < 0.0001). Fourteen tick species including engorged females from three genera were found on camels and co-herded livestock with varying prevalence (Supplementary Table 4). Sheep were infested only with *Rhipicephalus pravus* ticks. Three species were collected from goats (*R. pravus, H. dromedarii*, engorged *Rhipicephalus* spp. females) and five species from cows (*H. rufipes, A. gemma, H. dromedarii, A. variegatum* and *R. pulchellus*) (Fig. 1). We also observed that *A. gemma* was the most abundant tick on cattle but *H. rufipes* and *H. dromedarii* were the most prevalent species on camel. With our multiyear follow up ticks count on camels, we found that the prevalence of ticks on camels varied between years (Supplementary Fig. 1).

**Fig. 1.**
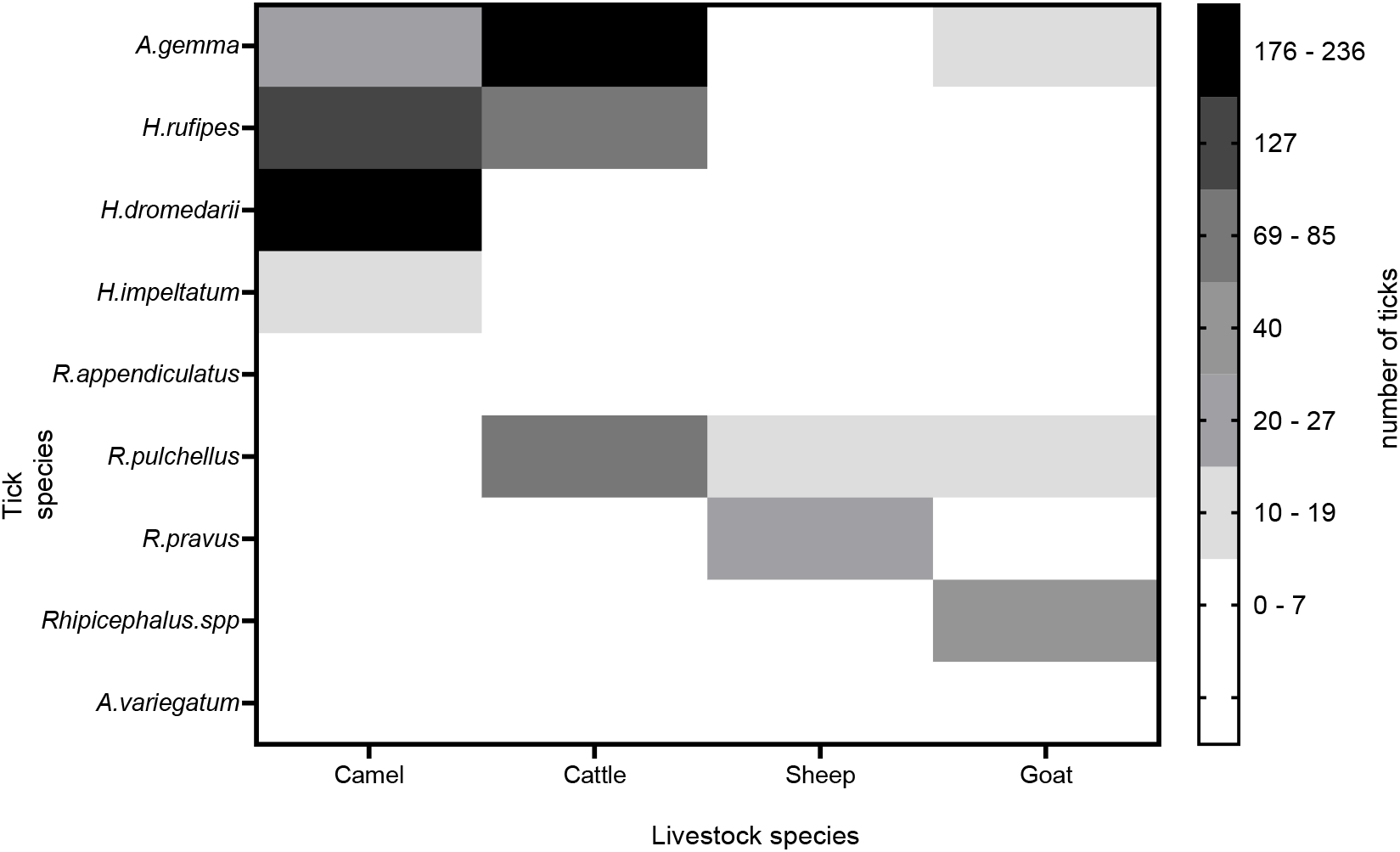
Heatmap showing tick density from different livestock hosts collected in 2019.

### Morphological Identification of Ticks

Since camel was the most preferred host (Fig.1) our further detailed study focused on camel-tick interaction. A total of 2494 hard tick species (Acari: Ixodidae) were collected from camel hosts from various ecologies. The ticks belong to three genera *Hyalomma* (74.5%), *Rhipicephalus* (15.4%), and *Amblyomma* (10.1%) collected from camels, from Marsabit and Samburu counties in Northern Kenya.

Based on morphology, 14 tick species in three genera were identified, namely, *Amblyomma gemma, Amblyomma lepidum, Amblyomma variegatum, Hyalomma dromedarii, Hyalomma impeltatum, Hyalomma marginatum, Hyalomma rufipes, Hyalomma truncatum, Rhipicephalus appendiculatus, Rhipicephalus camicasi, Rhipicephalus pravus, Rhipicephalus pulchellus, Rhipicephalus sanguineus*, and we also report occurrence of *Rhipicephalus simus* in the area, which was not reported previously (Fig.2). Detailed descriptions of the morphological features of the tick species are enlisted in Supplementary material 4.2. *H. rufipes* was the abundant species (27.7%) followed by *H. dromedarii* (25.8%); *R. simus* was the least abundant (0.03%).

### *Amblyomma gemma* Complex

We identified two novel *A. gemma* ticks which we designated as *Amblyomma gemma* variant whose postero-meridian stripes were narrow and not continuous (existence of bridges) despite the connection with the falciform stripe. Additionally, both ticks had narrow postero-accessory stripes while one had a distinct narrow cervical stripe. The lateral spots for the variants also varied as one had two lateral spots while the other had three seemingly as the definite *A*.*gemma*. Despite having pale rings, the legs of one of the variants were relatively darker in color (Fig. 3).

*Amblyomma gemma* ticks were identified by the flat slightly convex eyes close to the margin of the scutum; the localization of the primary punctation distribution on the scutum; the pink to orange enamel color; legs colored with pale rings; the short internal and medium-sized external spur lengths of coxae 1 and the small to medium punctations around the ocular region. Male *A. gemma* was distinguished by the presence of partial enamel ornamentation on 6 of the 11 festoons and the broad posteromeridian stripe; the elongate mesial area of enamel ornamentation and large and complex lateral median areas of the conscutum. The adult female *A. gemma* were identified by the straight scutum sides; broad scutum posterior angle; the large mesial area and the elongate and complex lateral areas of enamel ornamentation of the scutum.

### Molecular confirmation of tick species dynamics

Phylogenetic analyses were conducted on ticks collected from camels and co-herding livestock in northern Kenya employing molecular markers. We demonstrate the efficacy of 12S ribosomal DNA (rDNA), 16S rDNA, the nuclear ribosomal internal transcribed spacer 2 (ITS2), and cytochrome oxidase subunit 1 (CO1) genetic markers for DNA barcoding of ticks. These markers, along with existing morphological taxonomic keys, allow us to uncover the diversity of tick species (Fig. 4A-D). Our integrated approach, combining morphology generated from high resolution images with multiple molecular markers will facilitate the correct taxonomic identification of these arthropod vectors.

### Ticks’ diversity across various ecologies

Here, we describe the diversity of adult ticks in fourteen localities. We found significant differences in the diversity of tick species between sites (P<0.05). The Parkichon Fora and Marti Dorop sites recorded the highest Shannon diversity index (H′=2.2) with 9 out of the 14 recorded tick species, followed by Ngurunit (H′=2.1) while El-gade and Aiguman-Kalacha had the lowest (H′=1.1) with only 3 out of the 14 tick species encountered (Fig. 5).

### Infectious agents in Camels and associated *H. rufipes* ticks

To decipher the potential vectorial capacity of *H. rufipes* (the most prevalent tick on camel) regarding the pathogens it harbors and reservoir role of camel, we studied host-vector pathogen network by analyzing 299 *H. rufipes* ticks and all the camel blood samples collected for occurrence of tick-borne pathogens and other vector-borne pathogens such as trypanosomes. From our pathogen analysis between *H. rufipes* and camels, we found that some pathogens are shared between *H. rufipes* and camel. For instance, in *H. rufipes* ticks we found 4 pathogens including *Anaplasma spp. Ca*. Ehrlichia regneryi, *Rickettsia aeschlimannii* and *Coxiella burnetii* (Fig. 6A). However, we observed different pathogens dynamics in camels, for instance *Ca*. Anaplasma camelii and *Trypanosoma evansi* were the most prevalent pathogens in camels, but these were not detected in ticks. Moreover, *Rickettsia aeschlimannii* was abundant in ticks but was not detected in camels (Fig. 6B). The sequences of the various pathogens distinctly clustered into four different pathogens and grouped with known respective reference sequence (Fig. 6C).

### Spatial diversity of tick species on various Camels body

We asked if **t**he spatial variation in temperature, humidity, and microbes may create different ecological niches within a given host where ticks may thrive. Thus, we investigated the spatial distribution of tick species on various anatomical parts of camel. We found ticks are widely distributed across various body parts of camel. However, the nostril and anal region were the most infested sites, with nostril being the most preferred site by diverse species of ticks. Up to 50 ticks belonging to six different tick species were encountered per nostril in some camels (Fig.7A-B). The ubiquitous infested sites across the sampled livestock hosts were the ear and tail except for goats. The anal region of camels and cattle was infested mostly by *H. rufipes* and *R. pulchellus*. Other attachment sites on cattle included the dewlap, udder, the belly and along the neck while a few *A. gemma* were found on the male goat scrotum. The location of the tick attachment site is of clinical and tick-host interaction importance because it can allow for ticks to be rapidly discovered and removed, curtailing their ability to transmit pathogens. Additionally, the bite site may be pivotal in establishing disease outcomes.

### Camel-derived odours are attractive to *H. rufipes*

Having identified camel as the most preferred host and nostril as the most preferred predilection sites. We asked if *H*.*rufipes* are exploiting semio-chemicals to find their camel host. First, we analyzed if there are breath-specific odors that attract ticks to the camel nostril (Fig. 8A). We identified signature odors of camel breath as xylene isomers, 1,2,4-trimethylbenzene, cymene complex, mesitylene, dodecane, and tridecane. On the other hand, the body signature odors were, naphthalene, guaiacol, p-cresol, decanal, acetophenone, pinene, nonanal, and 1-octen-3-ol (Fig. 8A) (raw data obtained from ^15^, under a Creative Commons Attribution 4.0 International License) and the attractivity of body and breath signature odours was evaluated in a custom-made walking tunnel bioassay using individual compound against solvent control (Fig. 8B) using lab reared adult male and female *H. rufipes*. We found that *H. rufipes* was attracted to breath specific odor, specifically to dodecane t=4.34, df=9, P=0.002, p-cymene, t=3.72, df=9, P=0.005 and camel body odor 1-Octen-3-ol, t=4.4, df=9, P=0.002 which is a general odor present in various biological samples. The attraction for dodecane and 1-octen-3-ol was stronger. Interestingly, many of the other body and breath signature odors were neutral (Fig 8C).

## Discussion

Tick-host interactions influence the likelihood of pathogen acquisition, multiplication and transmission. In our ticks-pathogens-host interaction using high-resolution images coupled with four mitochondrial DNA (mtDNA) molecular markers, we revealed 14 tick species complexes feeding on various livestock. We characterized unique and shared pathogens between ticks and camel hosts. Finally, we elucidated chemical communication between *H. rufipes* and camels. Our study on the relationships between tick vectors-host-pathogens interaction is relevant to understand adaptation, evolution and to discover new targets for the development of innovative strategies to control vector-borne diseases.

Besides the fourteen tick species, we also identified two previously unreported, novel *Amblyomma gemma* variants. The sympatric nature of the populations of *A. gemma* and *A. variegatum* could be one of the reasons for the occurrence of the interspecific hybrid, thus leading to the reported *A. gemma* variants identified in the present study. Hybridization between ticks that are closely related in their sympatric zones is not exceptional^16^. The *A. gemma* variants had a few mismatches with the sequences of some of our samples and the previously reported *A. gemma* ^17^ collected at different locations even from northern Kenya. Consequently, the phylogenetic analyses indicated distinct clustering of the *A. gemma* variants across the genetic markers which were successfully amplified (16S, 12S and CO1) for molecular identification of the ticks.

Our data also reveals the occurrence of an undocumented Rhipicephaline tick species, *Rhipicephalus simus* which has not been reported from other studies in northern Kenya ^17,18^. The various techniques used in this study may have contributed to the identification, which signifies the importance of rigorous application of various taxonomic techniques. Moreover, our report of *Rhipicephalus pravus* corroborates the work by Dolan et al.^18^, conducted four decades ago from a camel herd in the Ngurunit area which was one of our study sites. The molecular markers mtDNA (CO1, 12S, 16S) and nuclear DNA, particularly, nrRNA gene (ITS2) were selective, thus some tick species were amplified using two or three markers but not the others. The ticks’ selective gene fragment amplification may be attributable to arbitrary or particular oligonucleotide primer designs. We suggest that the countermeasure of the discriminatory amplification phenomenon using three to four genetic markers and morphological analyses for certainty in species identification^19–21^.

In previous studies *Hyalomma dromedarii* was the most common tick species found on camels, accounting for 90% of infestations ^17,22–27^. However, we observed a considerable abundance of *H. rufipes* on camels. Changes or diversification in host species are tightly associated with organism genome evolution and species differentiation for host-obligatory organisms. For instance, there is a positive correlation of the increase in host switching with butterfly diversification^28^. In light of host switching, *H. rufipes* is mostly associated with cattle infestations^29^. The high infestation of *H. rufipes* on camels possibly as opportunistic hosts or as an evolutionary attribute that needs further investigation. Similarly, another instance is the collection of *H. dromedarii* from other hosts in the presence of their preferred hosts demonstrating ticks are not host specific the same as other blood-feeding arthropods relatives, insects. *H. dromedarii* has not been reported in other wildlife ecologies and non-arid and semi-arid lands (ASALs) in Kenya^30–33^. However, Ogola et al.^34^ demonstrated the importance of host availability despite optimum climatic conditions in an arid region where *H. dromedarii* was not found due to the absence of camels.

Several studies have indicated the predilection of *Hyalomma dromedarii* ticks as the only species that attach in the camel nostrils ^18,22,35^. Similarly, our spatial distribution map of ticks on various camel bodies confirms the proclivity of this tick species to attach to the camel’s nostrils. However, from our findings up to 50 ticks belonging to six different tick species were found in a single nostril, we also found, in addition to *H. dromedarii, H. rufipes, H. impeltatum, H. truncatum, A. gemma*, and *R. pravus* demonstrating camel nostril is the preferred predilection site of various tick species. However, there are species-specific and life stage dependent preferences for specific location of attachment especially in human-preferring ticks. For instance, on humans, *Dermacentor variabilis* preferentially bites the head and neck, while *Amblyomma americanum* prefers the thighs, groin, and abdomen, and *Ixodes scapularis* are found across the body ^2^. However, *Ixodes scapularis* showed a significant life stage difference with adults preferring the head, midsection, and groin, while nymphs/larvae preferred the extremities ^2^. Interestingly, infection resulted in a significant change in attachment site^2^ and infected tick attachment sites also determined the outcome of the infection, for instance, bite at the bite-site of single tick bites that resulted in infection with the Tick-Borne Encephalitis virus (TBEV) was associated with lethal outcomes if the bites were located on the head, neck, arms or axilla, while less lethality was associated with bites to the lower limbs and groin ^36^.

Relative to our findings, the reason behind various tick preferences for camel nostrils and the absence of ticks from the other three livestock nostrils is not fully understood and needs further investigation. This information is valuable for predicting the biting location of ticks. It is beneficial for the public to inspect several anatomical locations for ticks, with particular attention to the camel nose, which is often overlooked by pastoralists due to its hidden location. This can then facilitate the quick removal of ticks to prevent possible pathogen introduction, and potentially reduce the transmission of tick-borne pathogens or pathogen testing of the tick for diagnostic considerations. It is important to describe camel nostril from its biochemical, microbes, temperature, and pH among other elements. features to find out why it is the preferred niche for various ticks to thrive.

We characterized several pathogens from dromedary camels and *H. rufipes*, and document qualitative difference of the infectious agents between camels and *H. rufipes. Candidatus* Anaplasma camelii was highly prevalent in camels (66.3%) followed by *T. evansi* (8.8%), albeit these were not found in *H. rufipes*, that may demonstrate *H. rufipes* as an inefficient vector for these pathogens. Thus, potentially, the presence of the disease-causing microorganisms in camel could be due to the occurrence of other vectors including biting flies like *Stomoxys calcitrans* and *Hippobosca camelina*^37–40^. Some of the pathogens identified in *H. rufipes* corroborate with the report of TBPs in the same tick species from camels in Northern Kenya ^17^; these include *Coxiella burnetii, Rickettsiae aeschlimannii, Candidatus Anaplasma camelii*, and *Candidatus Ehrlichia regneryi. C. burnetii* and *R. aeschlimannii* characterized in *H. rufipes* ticks are of zoonotic importance as they are most likely to infect farmers who domesticate animals. Generally, ticks are known to be the reservoirs of *C. burnetii*, a neglected zoonosis and the causative agent of Q fever. On the other front, explicit reservoir animals are goats, cows, and sheep ^41,42^. Interestingly, the study by ^42^ reported on the high seroprevalence of *C. burnetii* in female camels, a case which was crosslinked with a history of abortions in the animals. The presence of *C. burnetii* in *H. rufipes* but missing in camel may demonstrate camel cleared the pathogen by the time of blood collection but remains in *H. rufipes* or *H. rufipes* interacted with other host before camel from where it has picked the pathogens, as *Hyalomma* species are two-host ticks. The mismatch in some pathogen between camel and tick demonstrates the use of tick for disease diagnosis may not provide the whole story unless we know it is a competent vector of the pathogen of interest and needs more detailed study from various host and ecologies.

The volatilome comparison between breath and body from the same camel showed each body part has its signature odors that are unique and shared but varies in abundance^15^. After identifying camel, as the most preferred host and *H. rufipes* the most abundant tick feeding on camel, we asked how they communicate and we showed their communication involves volatile organic compounds (VOCs) as *H. rufipes* exhibited a strong behavioral response to selected compounds, a response that has been selected over evolutionary time. From 14 signature compounds tested, dodecane and p-cymene were novel attractants to *H. rufipes*. Another attractant, 1-octen -3-ol, is a shared odor between various livestock and has been reported as an attractant to different tick species^43–45^. Similarly, other tick species are attracted to a few livestock-derived odors^43–45^. *A. variegatum* was found to be a generalist attracted to several odorants^46^ demonstrating tick species specific dependent bioactivity of livestock derived odorants.

The development of chemical ecology-based approach for tick management is timely for creating innovative strategies to control ticks and limit the spread of tick-borne diseases, especially as the nostril is a sensitive tissue for chemical application besides multiple acaricides resistance by various ticks. The localization of ticks in the camel nostrils is detrimental to the health of the animal due to the discomfort to the animal ^18^. However, despite other vertebrate hosts’ production of CO_2_ ^47^, the inclination of several species of ticks only to the camel’s nose is an interesting phenomenon that needs to be unravelled. Due to its occurrence in relatively every climatic region, *H. rufipes* has a wide distribution in Africa from desert to rainforest regions ^29,48,49^, thus understanding its adaptation mechanism both to various host, vectorial capacity and ecologies is the research of interest. The possible ecological effect of livestock diversification due to climate change, for instance, the shift to camel as a climate adaptation strategy concerning vector dynamics and pathogens transmission needs to be investigated in detail in the future.

In summary, fourteen species including two novel *Amblyomma gemma* variants of ticks feeding on camel were documented that varies in their diversity from place to place. Camels are the most preferred livestock by several tick species as compared to other co-herded livestock. Camel nostrils are the most preferred predilection sites from all anatomical regions. Finally, we have dissected the chemical communication of *H. rufipes* and camel by identifying potential attractants dodecane, 1-octen-3-ol and p-Cymene that may have potential in reducing the transmission of tick-borne pathogens. Our results provide insight about ticks-pathogens-host interaction, adaptation, reservoir and vectorial capacity and disease transmission dynamics.

## Materials and Methods

### Ethical Approval

This study was conducted within the framework of epidemiological surveillance activities at International Centre of Insect Physiology and Ecology (*icipe*), and the experimental guidelines and procedures were approved by *icipe*’s Institutional Animal Care and Use Committee (IACUC), reference number: *Icipe*ACUC2018-003-2023rev, and ethics approval obtained from the Ethics Review Committee of Pwani University (ERC/EXT/002/2020E). The ticks were collected from animals following authorization by the livestock owners after the objectives of the study were explained to them.

### Study Area

The study was conducted at Ngurunit, Moti, Dabala Fachana, Misa/Dabel, Bales-Bura, Aiguman-Kalacha, El-gade, Dokatu, Segel, and Malgis Lagha locations in Marsabit county, and Marti Dorop, Mpagas, Lependera, Parkichon Fora in Samburu County in Northern Kenya, where ticks were collected from camels and co-herding animals kept by livestock farmers (Fig.9). Marsabit county covers an area of 70,961.2 km^2^ and is located between latitudes 02° 45’ and 04° 27’ North and longitudes 37° 57’ and 39° 21’ East. The study area is climatically characterized as an arid and semi-arid ecology with minimum and maximum temperatures varying between 15°C and 25°C^50^. Rainfall is highly variable and erratic with an annual range of 200-1000mm, increasing as the altitudes rises^51^. Camel density in the area is high and contact between herds is high and regular. The lactation and reproduction status of the animals provide ground for the exchange of animals by the farmers between herds^52^. Other livestock kept in the area include cattle, sheep, goats, and donkeys. The animals are kept in homesteads referred to as “*bomas*”, which are enclosures usually fenced off with twigs; they include mud-walled housing and livestock holding areas. As a pastoral community, some farmers migrate from place to place in search of pastures and water resources and settle in camps (temporary settlements).

### Ticks’ Sampling and processing

Ticks were collected from livestock at the *bomas* in a cross-sectional study design in March 2019 (dry season), May 2020 (wet season), February 2021 (dry season), and June 2022 (wet season). The sampling was done early in the morning (0600-0900 hrs.). Ticks were removed from the animals using a pair of forceps and preserved in 99% absolute ethanol in 2ml Eppendorf tubes. The tubes were labelled according to the collection site, host species, attachment site on the host, and date of collection. The ticks were collected from different predilection sites of the animals which included the eye, ear, nose, body trunk, belly, udder, tail, and anal region. The collection tubes were frozen in liquid nitrogen in the field before transportation for analysis at *icipe*’s ML-EID laboratory where they were stored in -80°C freezers until further processing.

### Assessing ticks’ diversity between co-herding livestock

To study the ticks’ dynamics among co-herding livestock we selected only those households from the Ngurunit site that owned four types of livestock (camel, cattle, goat and sheep). Furthermore, due to the arid and semi-arid nature of the area, the number of cattle is low but the other three are abundant, with the small ruminants being the most dominant followed by camels^40^. We selected 10 households that fulfilled our criteria, i.e., those households that have all four species of livestock, and from each household we selected all tick-infested livestock and numbered, using the lottery method tick infested animals were selected randomly and counted the total number of ticks in all parts of the body. The total number of domestic animals sampled were 30 camels, 25 cattle, 20 sheep, and 20 goats.

### Camel Blood sampling

Blood was only drawn from tick-infested camels from the jugular veins by venipuncture into 10 mL disodium salt of ethylene diamine tetraacetate (EDTA) vacutainer tubes (Plymouth, PLG, UK). The vacutainers were temporarily stored in a cold chain (∼4°C) until completion of sample collection in the morning, following which, the blood was later transferred into 2 mL cryovials that were kept in liquid nitrogen before transportation to the laboratories for further processing. This study was approved by the International Centre of Insect Physiology and Ecology’s Institutional Animal Care and Use Committee (IACUC) (*icipe*-IACUC ref no. IcipeACUC2018-003-2023) and the Ethics Review Committee of Pwani University (ERC/EXT/002/2020E). All methods were carried out in accordance with relevant guidelines and regulations. Pastoralists/farmers gave their informed consent for their animal sampling after explaining the objectives of the study.

### Morphological Identification of Ticks

Using taxonomic keys^29,53^, ticks were morphologically identified to species level and sexed under a stereomicroscope focusing on the scutum ornamentation, body conformation, anal shields, and mouthparts. High-resolution images (Fig. 2 and 3) of both dorsal and ventral views of the respective tick species were taken to highlight the key identification features.

**Fig. 2:**
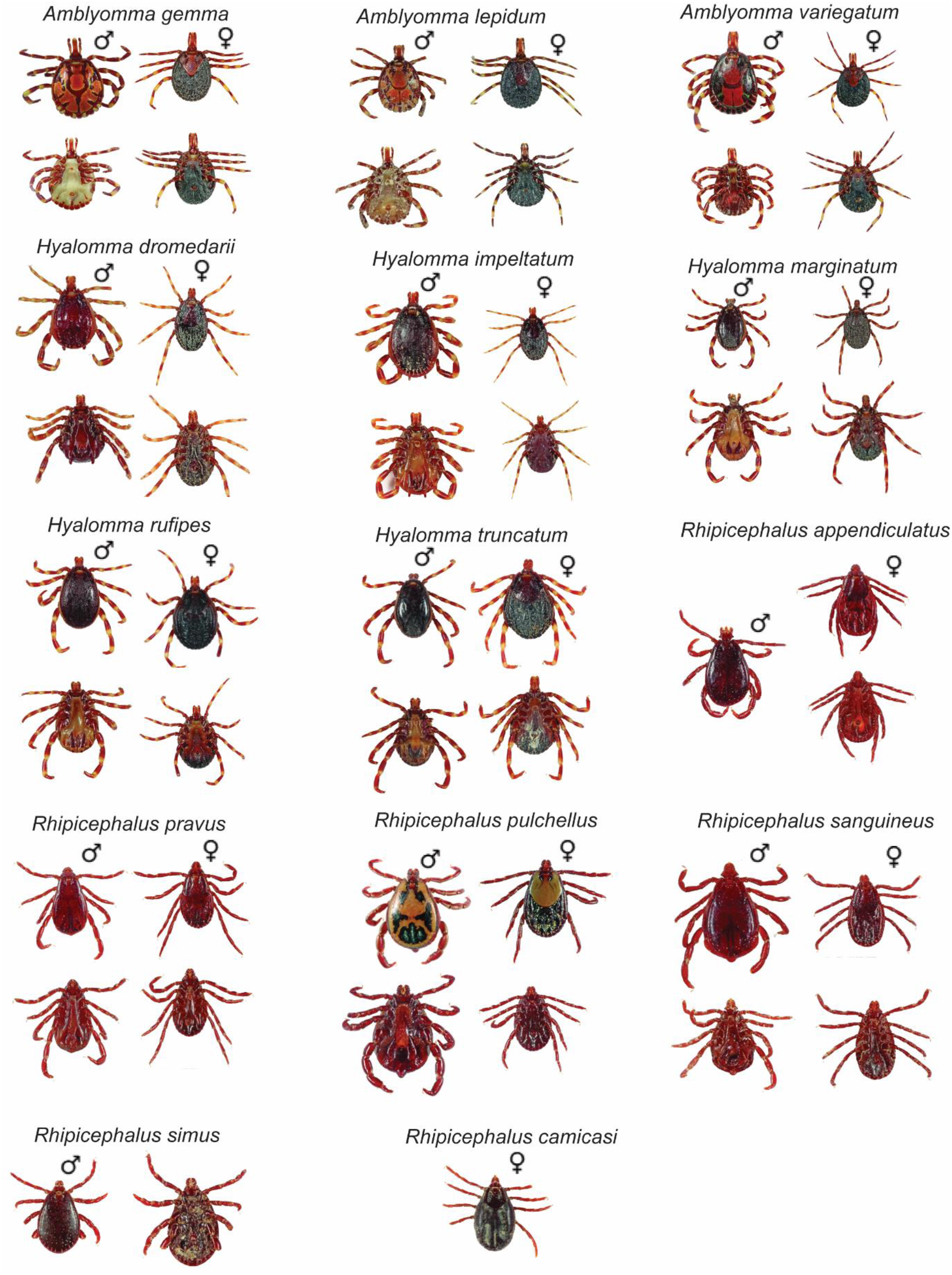
High-resolution photos of tick species identified from camels and co-herding livestock in Northern Kenya. ♂-male, ♀-female

**Fig. 3:**
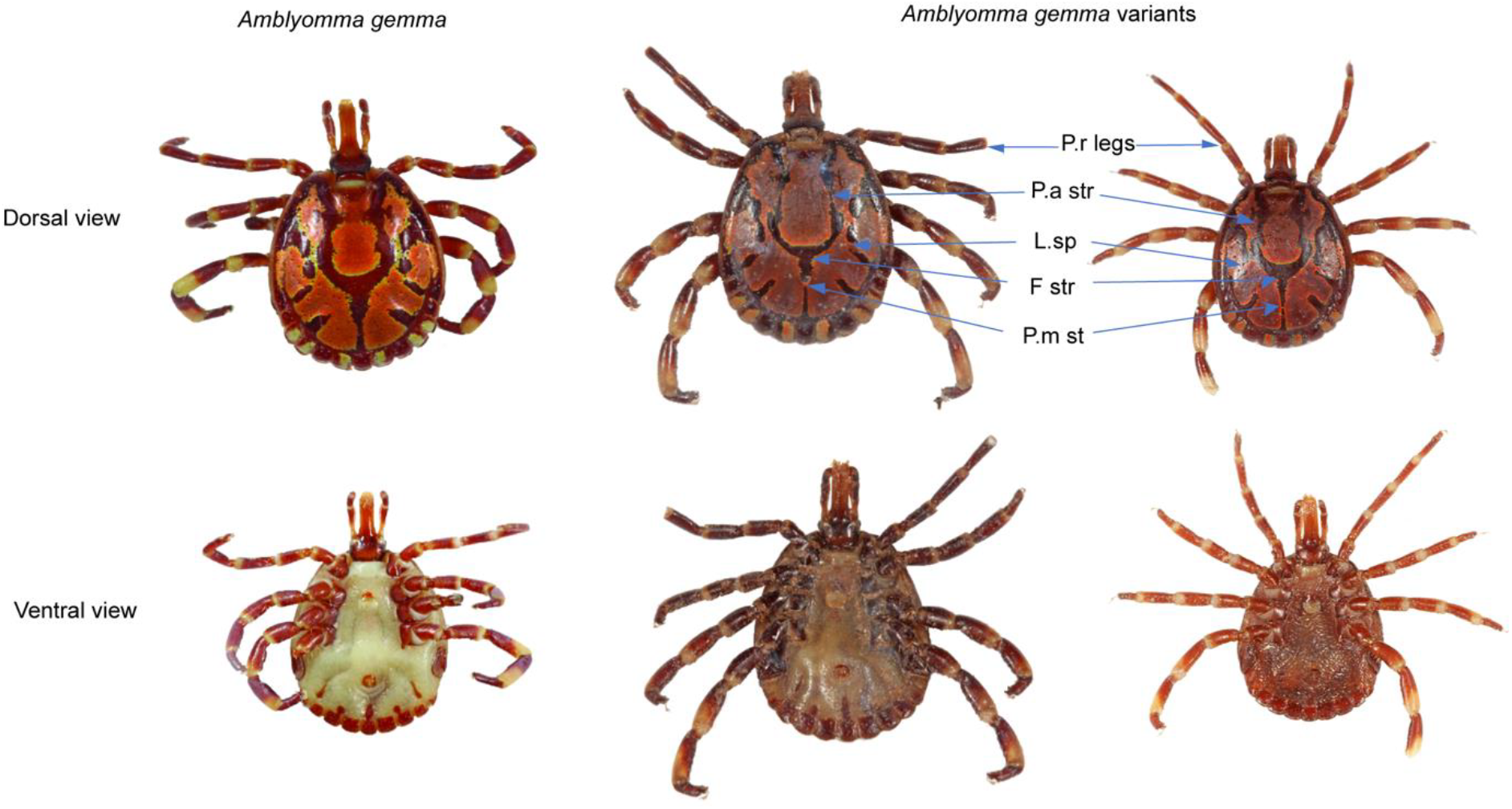
Differential morphological features of *Amblyomma gemma*. complex. P.r legs – pale-ringed legs; p.a str – postero-accessory stripes; l.sp – lateral spots; f strfalciform stripe; p.m st – postero-meridian stripes

### Molecular Characterization of Ticks and Tick-borne pathogens

To avoid PCR-inhibitors arising from contamination, exogenous DNA on the tick samples were removed by placing the ticks for one minute in a petri dish containing 1% sodium hypochlorite then rinsed for another minute in 1xPBS (Phosphate-buffered solution). The HotSHOT protocol was used in extracting the DNA from snippets of ticks’ legs for molecular identification of the tick species as described by^30,54^. DNA was extracted from (i) camel blood and (ii) *H. rufipes* ticks collected from camels using Isolate II Genomic DNA Kit using the genomic DNA extraction protocol according to manufacturers’ instructions^55^. After DNA extraction, the samples were subjected to conventional PCR for molecular characterization of the species of ticks and infectious agents in camels and *H. rufipes* ticks. A 20 μL PCR reaction mixture containing 4 μL HOT FIREPol® Blend Master Mix (Solis BioDyne, Tartu, Estonia), 1 μL of 10 μM reverse and forward primers (Supplementary Table 1)^56–65^, 4 μL of the template DNA, and 10 μL of nuclease free water was performed in a thermocycler (Applied Biosystems ProFlex PCR system). The cycling conditions for amplification of the respective target genes are indicated in the Supplementary Table 2.

### Agarose Gel Analysis, Amplicon Purification and Sequencing

10 μL of the amplicons were analyzed by running on 2% agarose gel at 100V for 1 hour before visualization on UV transilluminator. The remaining 10 μL of the PCR products for the positive samples were purified using EXOsapIT according to the manufacturer’s instructions^66^, and the cleaned products sent for Sanger sequencing at Macrogen Inc. (Amsterdam, Netherlands).

### Sequencing and Phylogenetic Analyses

All sequences were analyzed by trimming, editing and aligning using the Geneious Prime software (Biomatters Ltd., Auckland, New Zealand) v2023.0.4^67^ to generate consensus sequences. The consensus sequences were queried against the GenBank database using the Basic Local Alignment Search Tool (BLASTn) for identification of tick species and microorganisms. We aligned sequences from the present study against the reference sequences using Clustal Omega by grouping sequences by similarity as the alignment order to obtain consensus sequences. The sequences of the respective tick species and the pathogens identified in this study were deposited in the GenBank database; the accession numbers are shown in Supplementary Table 3. These sequences had >95% identity match with the existing sequences in GenBank. Further analyses were done to rigorously identify the ticks. Sequence alignments generated after aligning all the sequences using MAFFT plugin with default settings for each molecular marker were exported as Phylip. The sequences were subsequently cross-checked by aligning using ClustalW in MEGAX^68^. The exported Phylip files were used online to generate maximum likelihood phylogenetic trees using PhyML^69^ with 1000 replicates standard bootstrap analysis for 12S, 16S and ITS2. For CO1, a neighbor-joining (NJ) tree using the Kimura 2-parameter (K2P) distance metric (a standard model for analysis of DNA barcode data) as described by^70^ was constructed in MEGAX. The K2P model screens for contamination events, possible misidentifications, and other errors^71^. Tree visualizations (Fig. 4) were done using FigTree v. 1.4.4 (http://tree.bio.ed.ac.uk/software/figtree/). The percentage of replicate trees in which the associated taxa clustered together in the bootstrap test (1000 replicates) are shown next to the branches.

**Fig. 4:**
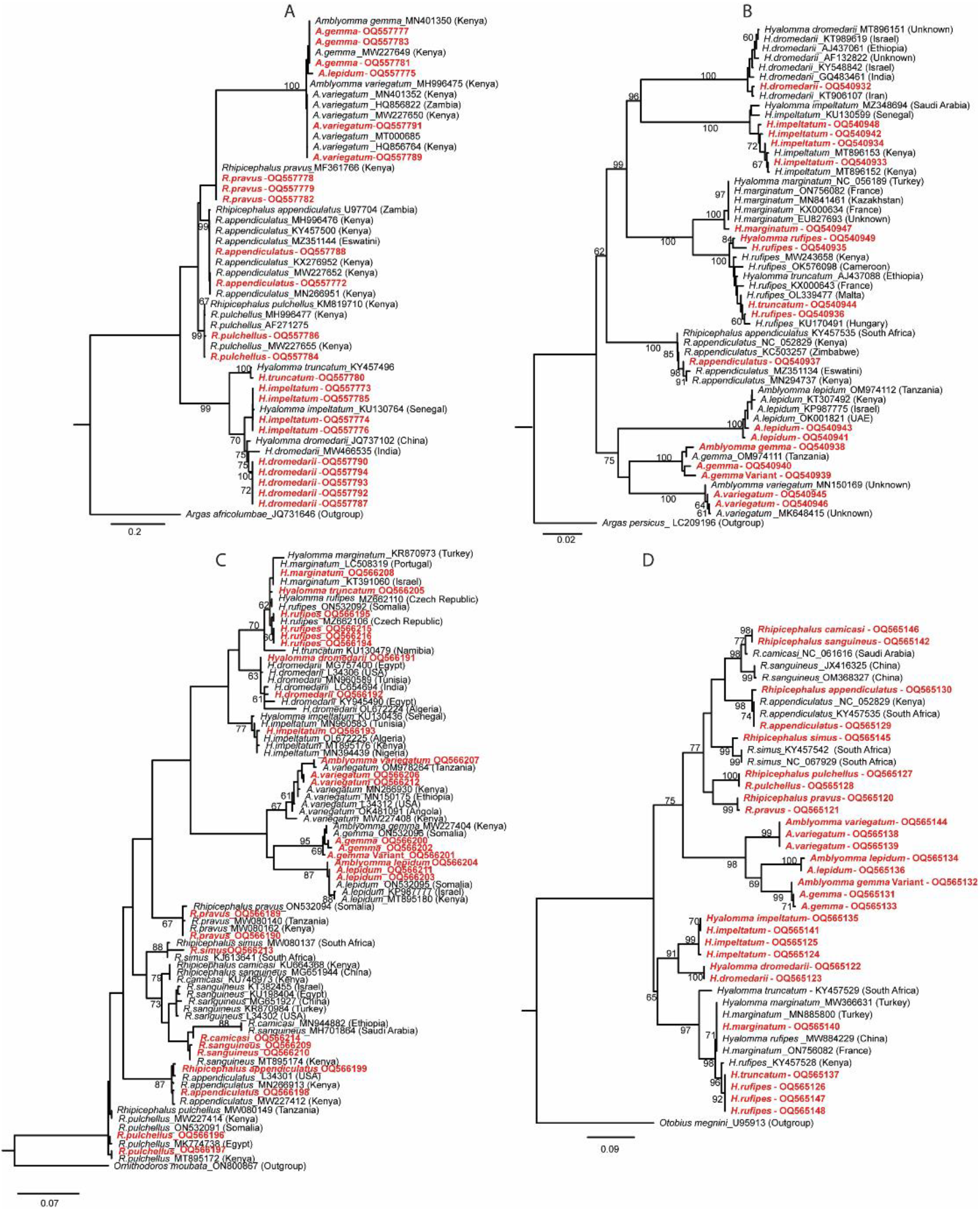
Phylogenetic analyses by maximum likelihood tree of **(A)** ITS2, **(B)** CO1, **(C)**16S rDNA, and **(D)** 12S rDNA. For CO1, neighbor-joining tree was applied. Soft ticks *Otobius megnini, Ornithodoros moubata, Argas africolumbae* and *Argas persicus* were used as outgroups for the respective molecular markers. Sequences obtained from this study are highlighted in red colored fonts.

**Fig. 5:**
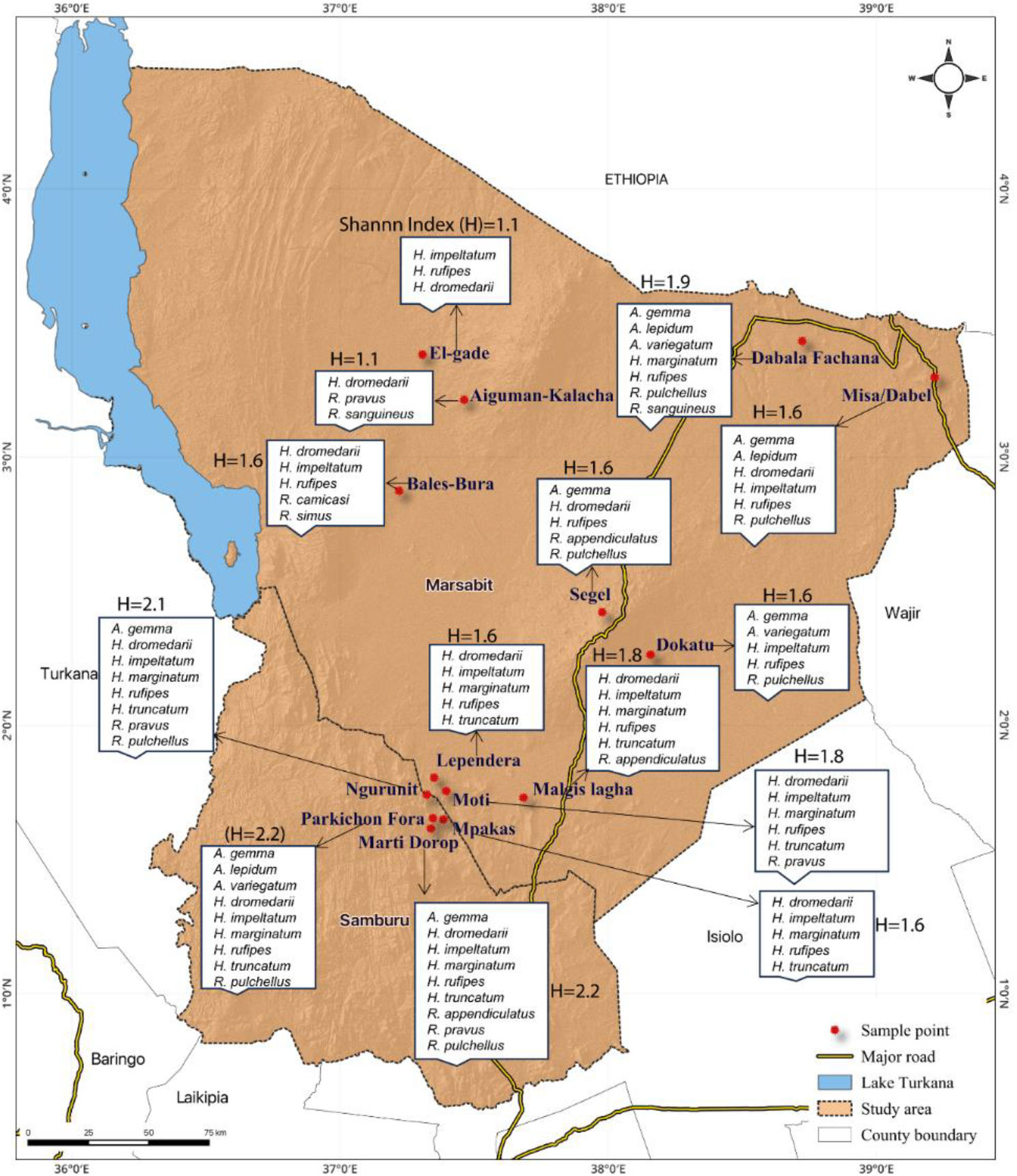
Locations of study sites and tick diversity. The various sampling sites were found to be different in ticks’ diversity, depicted by Shannon index (H).

**Fig. 6:**
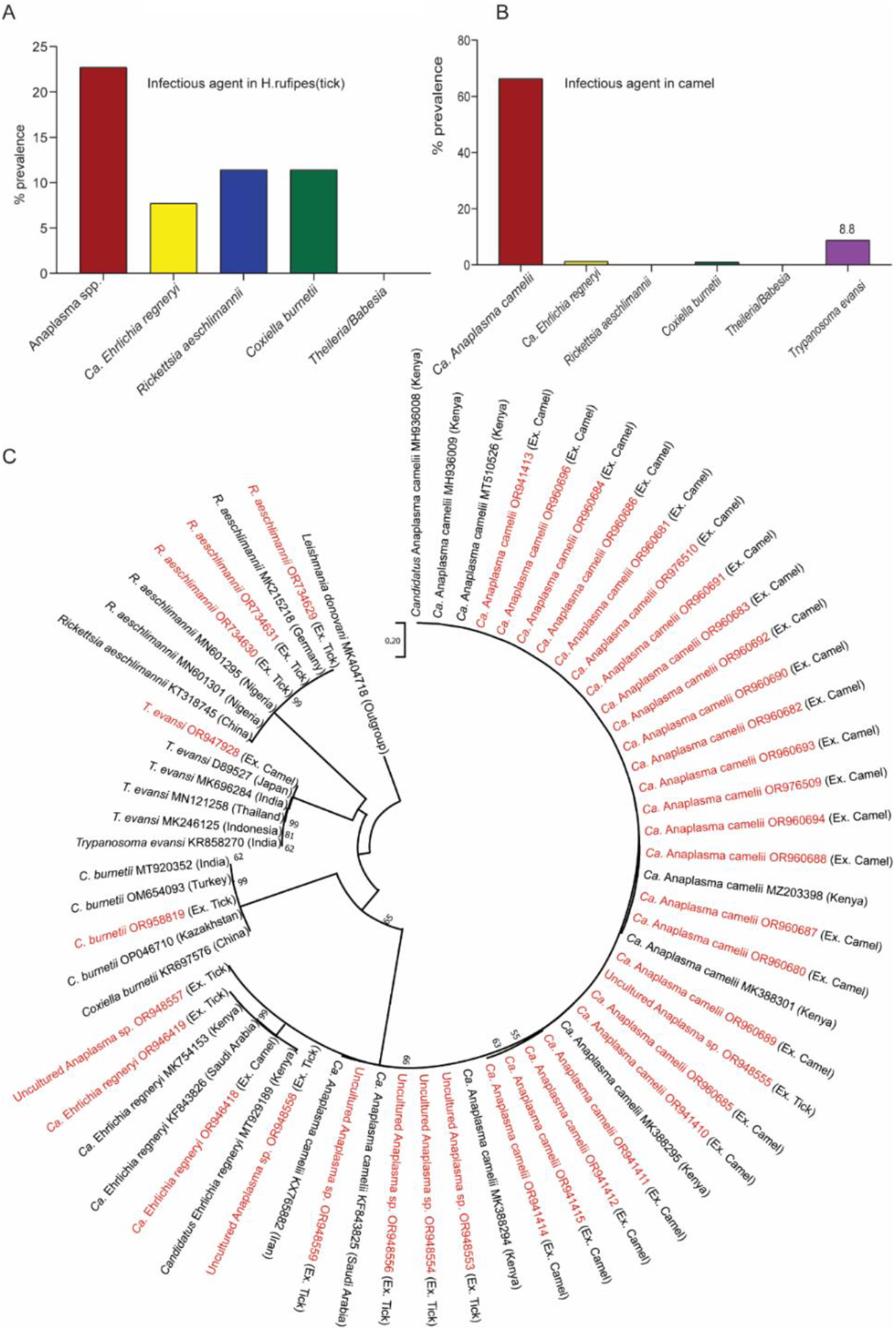
**(A)** Tick-borne pathogens characterized in *H. rufipes* ticks (n = 299); **(B)** Infectious agents identified in camels (n = 497) **(C)** Neighbor joining tree demonstrating the pathogens from camel and *H. rufipes* relatedness. Pathogens in red letters with their accession numbers are from this study.

**Fig. 7:**
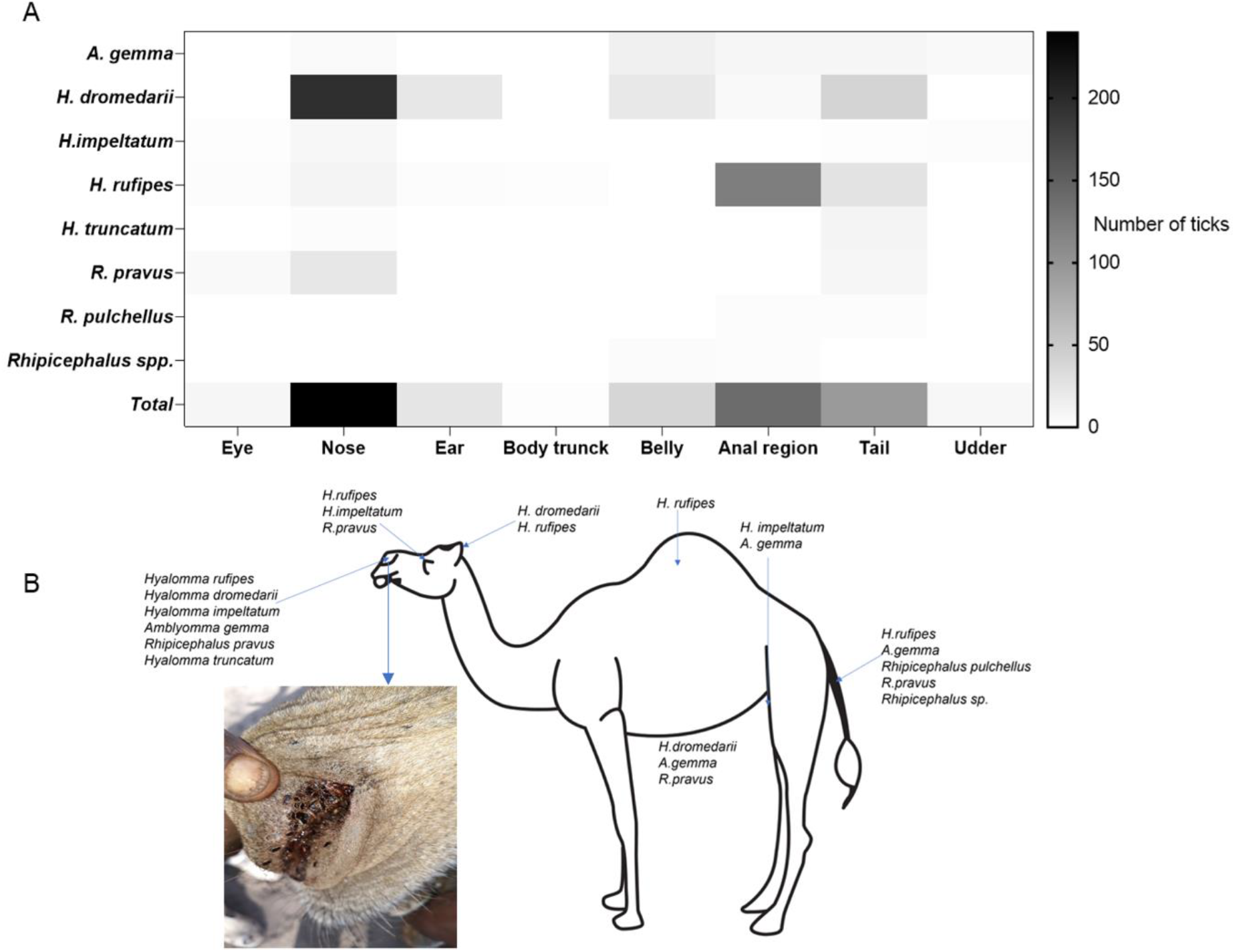
**(A)** Heat map showing the spatial distribution of tick species at different predilection sites on camel **(B).** Species distribution by predilection sites or tissue, inset shows highly infested camel nostril.

**Fig. 8.**
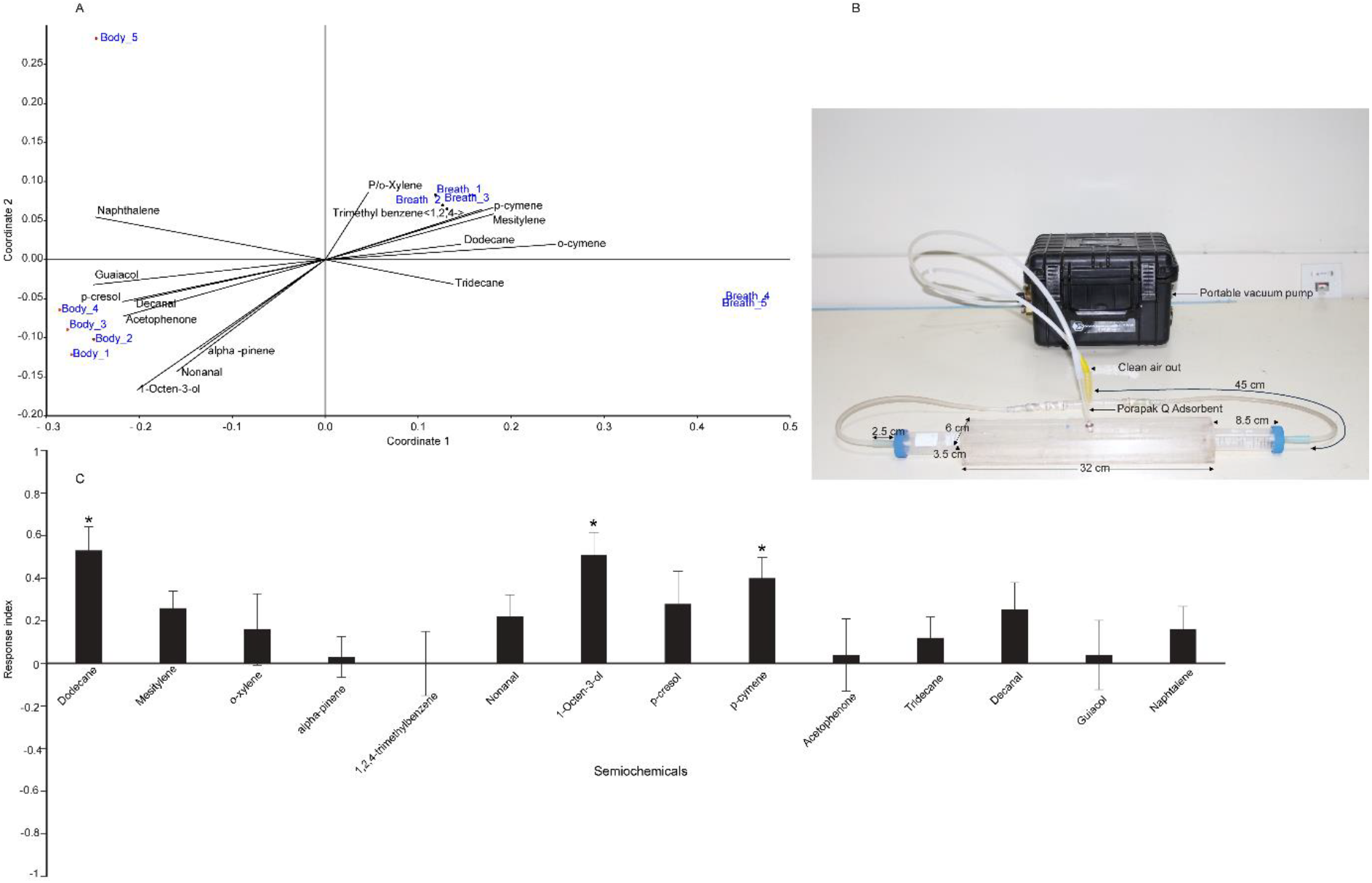
Camel-tick chemical communication. (A) Depicts signature odors of camel body and breath; (B) The custom-made walking bioassay; (C) The behavioural response of *H. rufipes* to various camel body and breath odors, *depicts a significant difference.

**Fig. 9:**
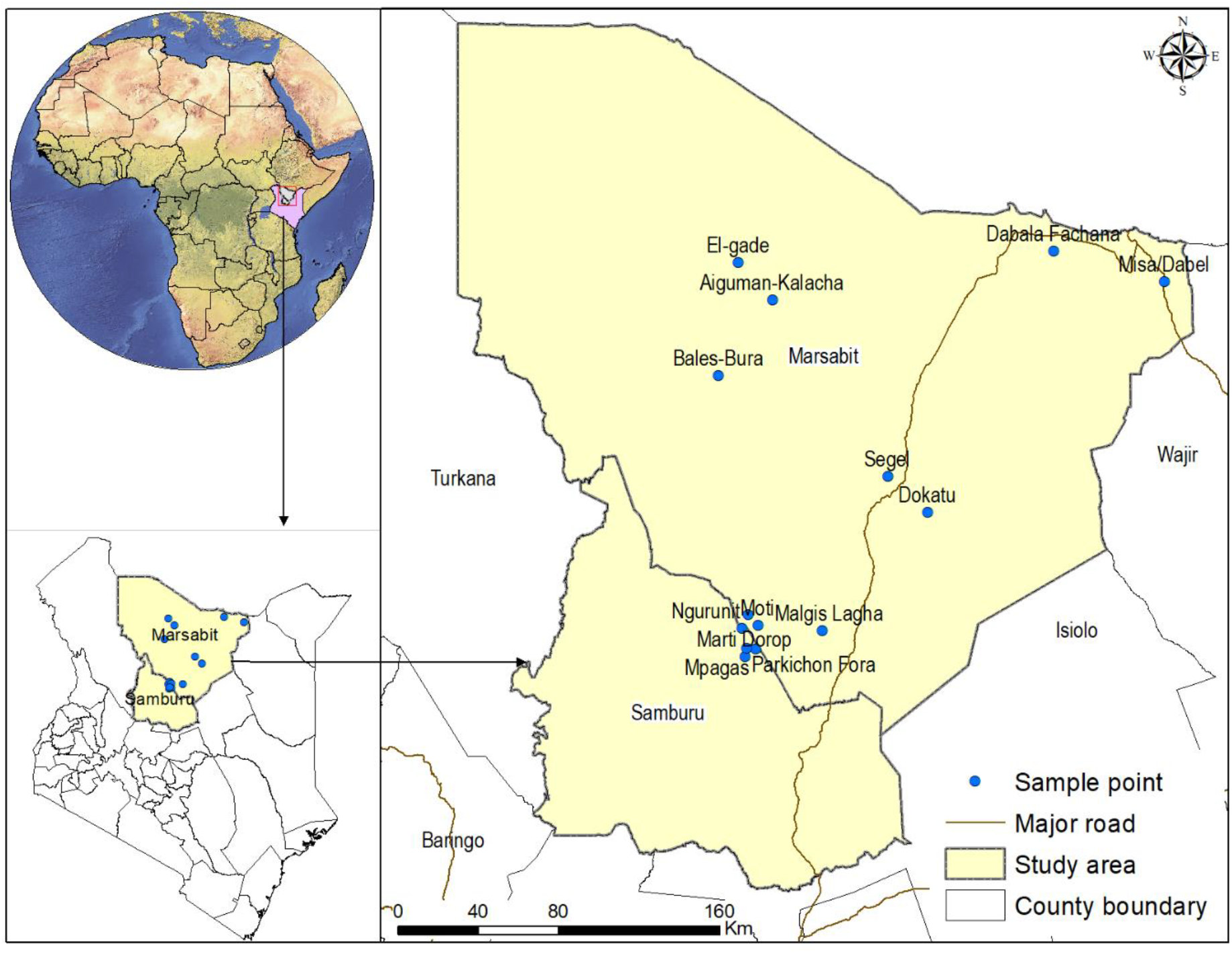
Map of the study area showing sites where ticks were collected from camels and co-herding livestock.

### Tick colony establishment

*Hyalomma rufipes* for the behavioral assay were obtained from a colony maintained at the *icipe* insectary. Adult *H. rufipes* were collected from naturally infested camels in Marsabit County, northern Kenya. Both the mature and immature stages parasitized successfully on New Zealand white breed rabbits. The rabbits were maintained at 50% relative humidity (RH), with temperatures ranging from 18 – 20°C, and exposure to natural daylight cycles. The rabbits were shaved on the back and a cotton fabric cloth glued on the shaved part and left for a day to firmly stick. After 24 hours, adult *H. rufipes* were introduced at the back of the animal, and closely monitored for successful feeding. During the non-feeding period, the engorged ticks were collected in separate cotton-plunged tubes which were maintained at an RH of 85 ± 5%. The RH was achieved by suspending the tubes in aluminium tins saturated sodium chloride solution. The tins were kept at 25 ± 1°C in a Sanyo MIR-153 incubator under photoperiods of 16:8 L:D. The longest period in the life cycle was the egg incubation, lasting close to 60 days. Hatched larvae were re-introduced on the host’s shaved back and placed on the glass tubes after feeding. Consequently, after the nymphs moulting, the number of adults were recorded. The life cycle of *H. rufipes* was averagely completed in 180 days: notably coinciding with the report by on the rearing of the same tick species under laboratory conditions^72^.

### Behavioral bioassay

Unfed *H. rufipes* adults were used in the behavioral assays. Treatment and control arms of the walking bioassay was loaded with 50 µl of odor and control on cotton roll. The odorants were diluted in hexane at 1µg/µl or 10^-3^ vl/vl and hexane was used as a control. Clean air at 85ml/min was pushed using porteable dynamic headspace odour collection vacuum pump (Sigma scientific USA). To avoid odor saturation at the middle of the tunnel we connected Porapak™ Type Q adsorbent at the tip of pulling tube of the pump. Five ticks of mixed sex were released in the middle and were given 5 minutes to make choice and decide, to avoid visual distraction the tunnel was covered with black cloth during the entire 5 minutes exploration and removed for tick choice counting. We counted the number of flies that had entered the treatment arm and control arm every 5 minutes. With this data, an attraction index (AI) was calculated as: 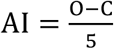, where O is the number of ticks that entered the treatment arm, or the odorant and C is the number of ticks that entered the control arm or hexane and 5 is the total number of ticks used per trial. The total number of ticks was limited to 5 to avoid aggregation and motivate navigation, when we use more than 5 ticks they were aggregating and no movement. The AI ranges from -1 (maximum avoidance) to 1 (maximum attraction). Zero denotes no choice between treatments. Fourteen camel derived odorants were tested and for each odorant the experiment was replicated 10 times.

### Statistics and Reproducibility

The statistical analyses were performed using various software such as R software version 4.0.3^73^ (R Core Team, 2020). The difference in tick prevalence between different livestock was analyzed using Generalised linear models with Poisson distribution using GraphPad Prism 10, using tick count data per livestock. Shannon diversity index (H) was used to determine the diversity of the tick species (richness and evenness) between the different sites using the number of ticks and the species collected per site. We applied NMDS Multivariate analysis using Bray-Curtis similarity measure to identify signature odors using Past version 3.02 from the headspace VOCs of the breath and body metabolic products. We employed the one-sample t-test, which is suitable for normally distributed data (confirmed by the Shapiro test with a p-value >0.05), to compare the attraction indices derived from the trap assay data with the theoretical mean of zero (0). For the analysis of the pathogen’s prevalence between ticks and camel to determine the sample size we used the formula 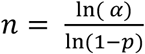 according to OIE manual for terrestrial animals 2012.

Based on our preliminary data, we identified several tick-borne pathogens including Candidatus *Anaplasma camelii* (8%), *Ehrlichia ruminantium* (3%), and *Coxiella burnetii* (1%) from ticks. We considered the low *C. burnetii* infection as our reference point for calculating the number of ticks to sample at 95% confidence limit. Using the formula 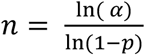, at α= 0.05, *P* = 0.01, the sample size n = -2.99/-0.01 = 299. The ticks were selected from all the ecological zones for the identification of TBPs.

## Acknowledgements

We thank John Ngiela and Victor Omondi for technical help in the field work; James Kabii, Dickens Ondifu and Dennis Getange for technical assistance in molecular studies; Raphael Mong’are and Juliet Onditi for designing the study sites’ map and Caroline Muya for handling administrative issues. Dr Steve Baleba for his useful comment on the original manuscript.

## Abbreviations

ML-EID: *icipe’*s Martin Luscher – Emerging infectious Diseases Laboratory

## Funding

This study was mainly supported by the Max Planck Institute-*icipe* partner group to MNG. Additional funding from German Ministry for Economic Cooperation and Development (BMZ) through the Deutsche Gesellschaft für Internationale Zusammenarbeit (GIZ)BMZ 81219442 Project Number: 16.7860.6-001.00. Norwegian Agency for Development Cooperation (NORAD), section for research, innovation, and higher education grant number RAF-3058 KEN-18/0005 (CAP-Africa) supported JOM. We also gratefully acknowledge the financial support for this research by the following organizations and agencies: The Swedish International Development Cooperation Agency (Sida); the Swiss Agency for Development and Cooperation (SDC); the Australian Centre for International Agricultural Research (ACIAR); the Norwegian Agency for Development Cooperation (Norad); the German Federal Ministry for Economic Cooperation and Development (BMZ); and the Government of the Republic of Kenya. The views expressed herein do not necessarily reflect the official opinion of the donors.”

## Contributions

J.O.M., M.N.G., and D.M. conceptualized and designed the research plan. M.N.G., P.N.N., F.A.O., and D.M supervised the work. PA assisted in tick collection and identification.

J.O.M. did the experiments and wrote the original draft. M.N.G., P.N.N., F.A.O., and D.M reviewed and edited the manuscript. All authors have read and agreed to the published version of the manuscript.

## Declarations Competing interest

The authors declare no competing interest following the reporting of the findings herein.

